# Radiation resistance in cervical cancer: An insight on mitochondrial associated membranes and its related proteins

**DOI:** 10.1101/2024.04.23.590692

**Authors:** Harini Lakshminarasimhan, Rashmi Santhoshkumar, Sweta Srivastava

## Abstract

Radiation therapy proves ineffective against radiation-resistant cancers, exacerbating the disease. The Mitochondrial Associated Membrane (MAM) forms through the tethering of the endoplasmic reticulum to the mitochondrial membrane. Under stress conditions such as radiation, mitochondrial function becomes impaired, prompting the recruitment of endoplasmic reticulum proteins for cell survival. Therefore, understanding the role of ER and mitochondria interactions is crucial for the development of inhibitors or drugs to sensitize cancer cells to radiation. In this study, we focused on ultrastructural alterations in radiation-resistant cells. The main structural changes observed included mitochondrial elongation, mitophagy, the presence of lipid droplets, and shortened ER segments. While MAM formation (the interaction between the ER and mitochondria) was observed in SiHa cells not exposed to radiation, it was disrupted in radiation-resistant cells. We analysed genes associated with mitochondrial elongation, mitophagy, lipid droplet formation, short ER fragments, and ER stress using a dataset from cervical cancer patients in The Cancer Genome Atlas (TCGA). Our analyses suggest that BNIP3L is implicated in therapy failure and tumour recurrence. In future, inhibiting BNIP3L could potentially sensitize resistant cells to radiation therapy.

## 1. Introduction

Radiation therapy is one of the major treatment modalities, alongside chemotherapy and surgery, for most cancers. Nearly 70% of patients receive radiation therapy in addition to surgery and chemotherapy [1,2]. Therapy failure not only impacts the quality of life but also imposes a financial burden on patients. Additionally, radiation therapy produces free radicals that damage DNA, inducing ROS and oxidative stress, ultimately leading to tumor cell death [1]. Radiation therapy also contributes to the development of radiation resistance in cancer cells, leading to therapy failure. This failure results in poor treatment outcomes, prognosis, cancer recurrence, and metastasis. Furthermore, radiation damages normal tissues surrounding cancer cells, triggering inflammation and bleeding. Radiation therapy employs high-energy photon radiation, such as X-rays and gamma (γ)-rays, to target cancer cells, inducing single-strand and double-strand breaks in DNA, inhibiting cell proliferation and division, and ultimately resulting in apoptosis and anoikis [2].

Cancer cells develop resistance to radiation through intrinsic or acquired mechanisms. Intrinsic resistance arises from changes in cancer genes or phenotypes within the cell, while acquired resistance is induced by the cancer microenvironment. Radiation resistance can be induced by a hypoxic environment, upregulation of DNA repair pathways, transitions in the epithelial-mesenchymal transition (EMT) pathway, activation of cancer stem cells, and inactivation of apoptotic pathways [1,3]. Recent studies have highlighted the importance of other cellular organelles besides DNA damage in radiation resistance [4].

Changes in mitochondrial size, shape, and mutations in mitochondrial DNA disrupt the normal physiological function of mitochondria, enhancing their adaptability to radiation. Radiation alters mitochondrial metabolism, ROS production, hypoxia, and mtDNA methylation status [4,5]. MtDNA is more sensitive to radiation damage than nuclear DNA. Mitochondria respond to radiation-induced DNA damage by increasing the copy number of mtDNA, and altering the expression levels of enzymes such as glutathione peroxidase, isocitrate dehydrogenase, manganese superoxide dismutase, and catalase to minimize radiation-induced damage. Additionally, mitochondrial membrane potential and metabolism are rewired to overcome radiation-induced damage, leading to resistance to radiation therapy [6].

While mitochondrial involvement in radiation resistance has received significant attention, its interaction with other organelles during radiation exposure remains to be fully understood. One crucial interaction is through the Mitochondrial Associated Membrane (MAM), which serves as the contact site between mitochondria and the endoplasmic reticulum. MAMs play an essential role in cell physiology and function, being a dynamic structure involved in lipid transport, calcium homeostasis, ROS production, autophagy, apoptosis, etc. [7]. However, the dynamics of MAM upon irradiation in cervical cancer cells have yet to be investigated. This paper aims to identify the changes in MAM induced by irradiation in radiation-resistant cervical cancer cells.

## 2. Materials & Methods

All the tissue wares were purchased from BioStar. Antibodies BNIP3L, TOMM20, MFN2 were from Santa cruz, ROCK2 and GAPDH antibodies were purchased from CST.

### 2.1. Cell Culture

SiHa cell lines utilized in this study were cultured in Dulbecco’s Modified Eagle Medium (DMEM) (Gibco) supplemented with 10% Fetal Bovine Serum (FBS) (Sigma), 50□U□ml^−01^ penicillin, and 50□mg□ml^−01^ streptomycin at 37□°C in a 5% CO_2_. Irradiation of cell lines was performed using a linear accelerator (LINAC). Cell viability assays were conducted using WST-1 reagent (Roche). Tissue wares were purchased from BioStar. Antibodies for BNIP3L, TOMM20, and MFN2 were obtained from Santa Cruz, while ROCK2 and GAPDH antibodies were from CST.

### 2.2. Development of Radiation-Resistant SiHa Cells

SiHa cells, cultured in 6-well dishes (BioStar), were irradiated with 6 Gy radiation [8] using a linear accelerator (LINAC). Following 7 to 14 days of radiation, these cells were again irradiated with 6 Gy radiation. This process was repeated until the cells received a total of 48 Gy radiation. All experiments were performed between the 5th and 14th day of radiation exposure.

### 2.3. Cell Survival Assay

1 × 10^4^ SiHa N and SiHa-Res cells were seeded in a 96-well plate. 5□μl of WST1 was added along with culture media and incubated for 30 min at 37□°C in a CO2 incubator. The absorbance of the plate was read at 450□nm using a microplate reader. Percentage of cell survival was calculated using the formula (Absorbance of treated cells/Absorbance of untreated control) x 100.

### 2.4. Clonogenic Assay

1 × 10^3^ of SiHaN and SiHa-Res cells were seeded in a 60mm dish and allowed to grow for 15 days in CO_2_ incubator. The plates were then washed, stained with methanol in 0.1% CBB and the size and number of colonies were noted.

### 2.5. CFSE Assay

SiHaN and SiHa-Res cells were grown on coverslips. The CFSE assay (Abcam, cell labeling kit) was performed according to the manufacturer’s instructions. Briefly, CFSE reagent was added, and coverslips were removed at 15 minutes, 48 hours, and 96 hours, then fixed and stained with DAPI. The coverslips were imaged using a fluorescence microscope (Leica).

### 2.6. qPCR

RNA isolation was performed using the TRIzol method according to the manufacturer’s protocol (Life Technologies, Invitrogen). M-MLV reverse transcriptase was used for cDNA conversion. Gene expression was analyzed by qPCR using Power SYBR green fast master mix and run on a 7500 Fast Real-Time PCR system by Applied Biosystems. Primer sequences used in the experiments are tabulated below.

Sequences of primers used

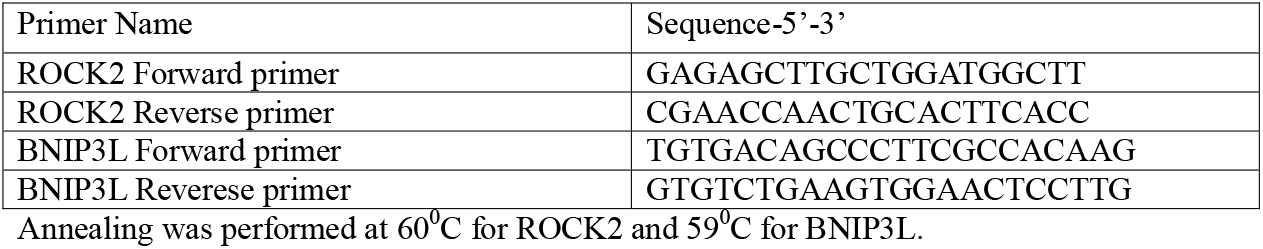

Annealing was performed at 60^0^C for ROCK2 and 59^0^C for BNIP3L.

### 2.7. Western Blot

SiHaN and SiHa-Res cells were lysed with RIPA cell lysis buffer and centrifuged. Electrophoresis was performed on the lysates, followed by transfer to a nitrocellulose membrane. After blocking, the membrane was probed for MFN2, ROCK2, BNIP3L, and Actin overnight. This was followed by secondary antibody incubation and detection with ECL chemiluminescence reagent (Invitrogen). Protein expression levels were represented as fold change relative to actin.

### 2.8. Immunofluorescence

Cells were grown on coverslips, fixed, permeabilized, and blocked. Primary antibodies were added and incubated overnight at 4°C. Secondary antibodies were used at a 1:500 dilution for 45□min, followed by staining with DAPI. The cells were then washed and mounted on a glass slide using Vector Shield (Vector Laboratories, INC).

### 2.9. Transmission Electron Microscopy

SiHaN and SiHa-Res cells in 60mm dishes were fixed with 2.5 % buffered glutaraldehyde, post fixed in osmium tetroxide, dehydrated using graded series of distilled ethanol (70%, 80%, 90%, 95% en-block with uranyl acetate and 100%), cleared using propylene oxide, infiltrate and finally embedded in araldite resin. Ultrathin sections of ∼80nm thickness were collected on copper grids and contrasted using saturated methanolic uranyl acetate and 0.2% lead citrate. Electron micrographs were captured using transmission electron microscope (TEM) JEM 1400-Plus (Jeol, Japan).

### 2.10. *In-Silico* Analysis

A cervical cancer dataset (GSE14404) [9] with 28 cancer patients (10 with radiotherapy failure and 16 with no evidence of disease after therapy) was utilized. Fold change in gene expression between no evidence of disease and failed cases was calculated. Overexpressed genes were further analyzed using UCSC Xena and GSCA portal for TCGA analysis.

### 2.11 Statistical Analysis

All experiments were performed in triplicates. Mean and standard deviations were computed. Student’s t-test was used for significance, with p□<□0.05.

## 3. Results

### 3.1 Radiation-Resistant Cells Exhibit Enhanced Survival, Proliferation and Clonogenic Ability

To understand the changes associated with radiation resistance, SiHa cells were serially radiated to create resistant cell line (SiHa-Res). Radiation resistance in SiHa-Res cells was confirmed through cell survival, proliferation and clonogenic assays. Irradiated SiHaN and SiHa-Res cells were subjected to a WST assay and clonogenic assay. Figure 1(A) illustrates that SiHa-Res cells have a survival rate of 86%, whereas SiHaN cells have a survival rate of 48% compared to the non-irradiated control SiHa cells. Clonogenic assay results depicted in Figures 1(B) show that SiHa-Res cells form nearly 5 times more colonies compared to SiHaN cells. Cell proliferation was assessed with CFSE assay. Cell proliferation was assessed using the CFSE assay. Figure 1(C) indicates that the uptake of CFSE by cells was highest at 48 hours. Subsequently, there was a decrease in fluorescent intensity in both SiHaN and SiHa-Res cells at 96 hours. However, at 96 hours, the fluorescent intensity of SiHa-Res cells was higher than that of SiHa N cells.

**Figure. 1.**
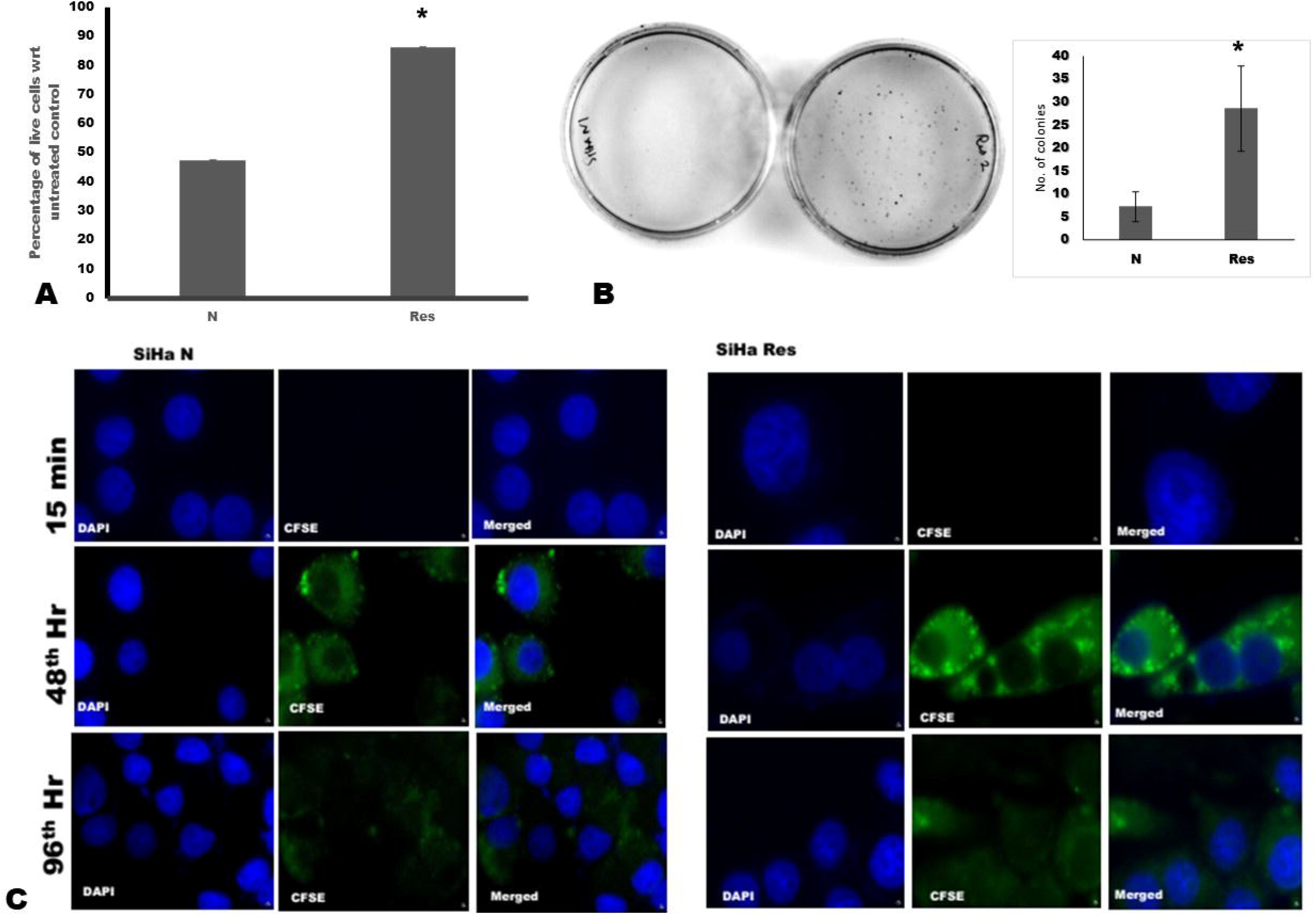
Cell survival & proliferation of SiHa normal & resistant cells. SiHa-Res cells (A) has enhanced survival and (B) clone formation than normal SiHa cells. (C) Proliferation of untreated SiHa cells and SiHa-Res cells with CFSE reagent. * indicates p< 0.05

Radiation-resistant cells were developed by repeated exposure of SiHa cells to 6 Gy radiation. The SiHa-Res cells used in this study acquired resistance to 48 Gy of radiation. Radiation resistance in SiHa cells was confirmed by WST and clonogenic assays. The percentage of live cells in Figure 1(A) indicates that SiHa cells were resistant to radiation, whereas the normal SiHa cells showed only 49% live cells compared to the non-irradiated control SiHa cells. Similarly, the clonogenic assay in Figure 1(B) indicates that SiHa-Res cells have an enhanced ability to form clones compared to SiHaN cells upon exposure to 6 Gy radiation. Figure 1 (C) illustrates the proliferation abilities of SiHa N and SiHa-Res cells. The proliferation of SiHa-Res cells appears to be lower compared to untreated SiHa cells.

### 3.2 Altered MAM Structure in Radiation-Resistant Cells

Ultrastructural studies were conducted to observe changes in cellular organelles in radiation-resistant cells. Transmission electron microscopy (TEM) images (Figure 2A) of SiHaN cells reveal close proximity between the endoplasmic reticulum (ER) and mitochondria, indicating MAM formation. Prominent MAM structures were also observed in Figure 2B, where the rough endoplasmic reticulum (RER) was adjacent to mitochondria. In SiHa-Res cells (Figure 2C), the formation of lipid droplets and mitochondrial structural changes were observed. Figure 2 (D) depicts a denser mitochondrion undergoing autophagy or mitophagy, while Figure 2(F) shows an elongated mitochondrion more than 500nm long. Interestingly, RER structures were also altered in radiation-resistant cells, as seen in Figure 2(B) for SiHaN cells and Figure 2(E) for SiHa-Res cells, where a shorter fragment of RER is visible. To confirm changes in MAM structure, the expression levels of MFN2, a MAM marker [10], were studied via western blot. Figure 2(G) compares the expression level of MFN2 in SiHaN and SiHa-Res cells, revealing a downregulation to 0.816-fold in SiHa-Res cells compared to SiHaN cells.

**Figure. 2.**
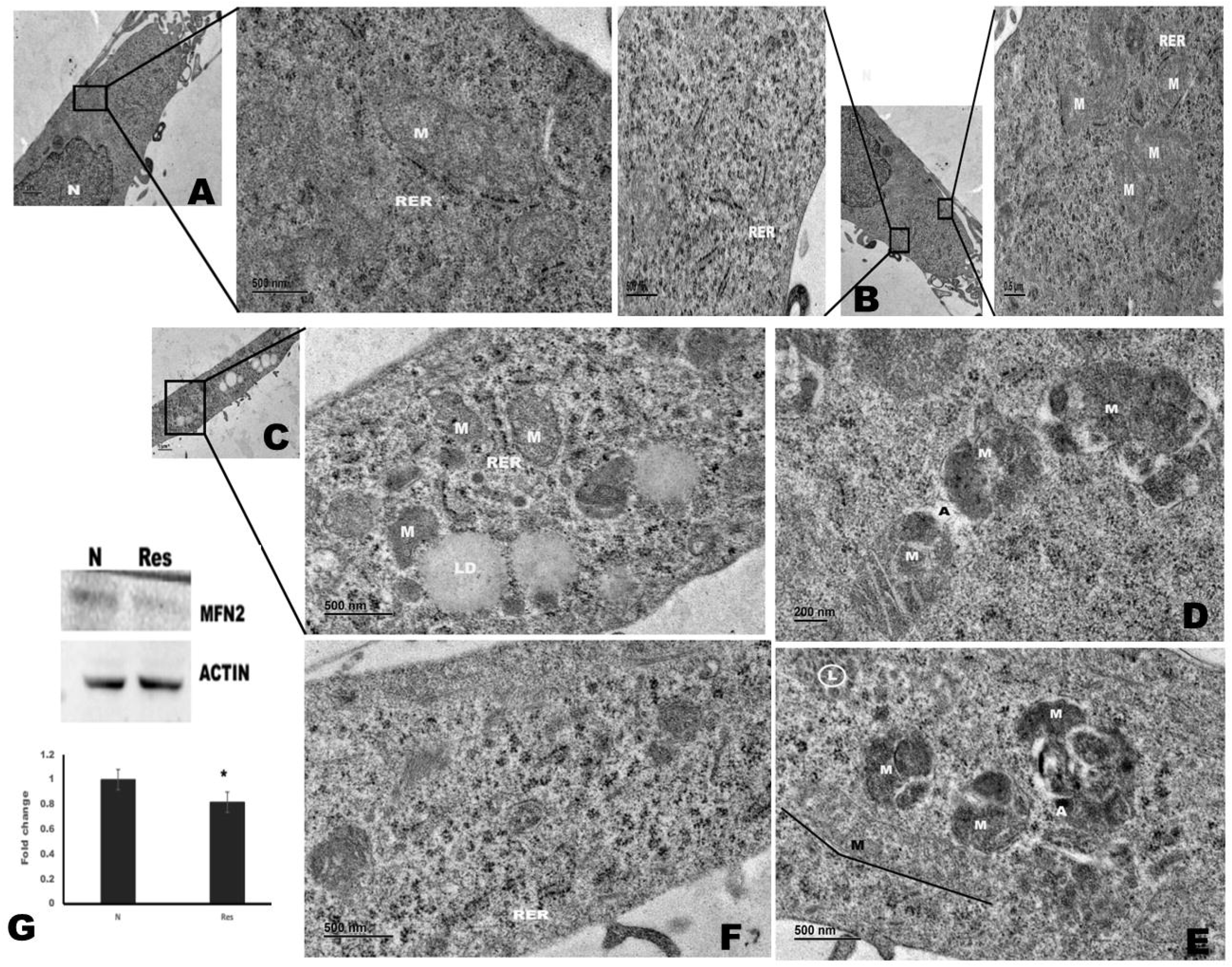
Disturbed MAM in Resistant cells. TEM of MAM in SiHaN cells with (A) RER in close proximity with mitochondria (2500x) and inset 5000x. (B) Long stretches of RER and RER in close association with mitochondria (2500x, inset both 8000x). TEM images of SiHa-Res cells. Resistant cells with (C) lipid droplets (2500x, inset 8000x) (D) mitophagy (10000x) (E) short fragments of RER (8000x) (F) elongated mitochondria which is more than 500 nm (8000x). (G) Western blot images and fold change graph of downregulation of MFN2 in SiHa-Res cells.

### 3.3 *In-Silico* Analysis for Target Gene Prediction

To identify target genes involved in radiation resistance, a patient dataset on radiotherapy failure and disease-free survival from GEO Accession ID GSE14404 [9] on cervical cancer was obtained. Genes that were overexpressed in failed sets (radiotherapy failure) compared to those with no evidence of disease (NED) were analyzed based on fold change. The gene list from the patient dataset was compared with genes from a Uniport search including terms such as “autophagy,” “lipid droplets,” “ER fragmentation,” “ER stress,” “mitochondrial elongation,” “mitochondrial dysfunction,” and “autophagy.” These genes were further grouped into categories such as lipid droplets, ER fragmentation, ER stress, mitochondrial elongation, mitochondrial dysfunction, and autophagy (Figure 3). From the patient dataset, 48 genes were associated with autophagy, 75 genes with ER stress, lipid droplets, and ER fragments, 29 genes with mitochondrial elongation and dysfunction, and 13 genes were shared between ER stress, lipid droplets, ER fragments, and autophagy. Additionally, 9 genes were common between ER stress-related, autophagy-related, and mitochondria-related sets. These genes were grouped and analyzed using FUNRICH [11]. The gene list was tabulated based on these categories (Figure 4). Genes responsible for cancer recurrence, such as ABCA1, BCAS3, BMP2, BNIP3L, CD84, CLU, CSTF2, GPR137B, GPR85, KLC1, NPY, PIK3R1, PNPLA8, and TRPC3, were obtained from the “TCGA PanCancer” study via the UCSC Xena portal [12], with tissue of origin selected as “CERVIX” (Table 1).

**Table 1.** Short listed genes from patient dataset for tumour recurrence in UCSC Xena.

| Shortlisted genes for tumour recurrence |
| --- |
| ABCA1, |
| BCAS3, |
| BMP2, |
| BNIP3L, |
| CD84, |
| CLU, |
| CSTF2, |
| GPR137B, |
| GPR85, |
| KLC1, |
| NPY, |
| PIK3R1, |
| PNPLA8, |
| TRPC3 |

**Figure 3.**
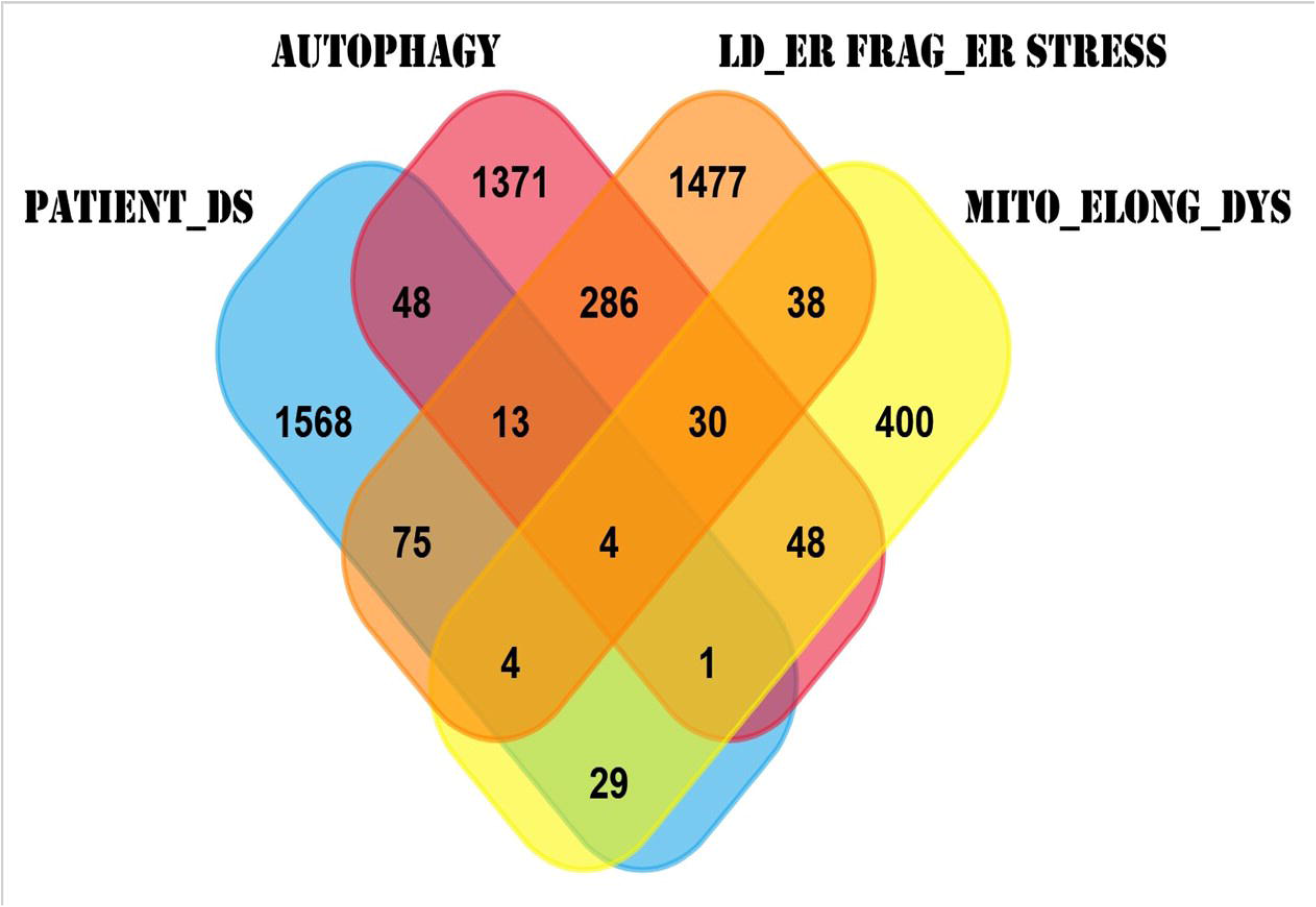
Venn diagram of common genes between autophagy, Mitochondrial elongation, mitochondrial dysregulation, ER stress, short ER fragments, lipid droplets and genes of cervical cancer radiation therapy failed patient dataset.

**Figure 4.**
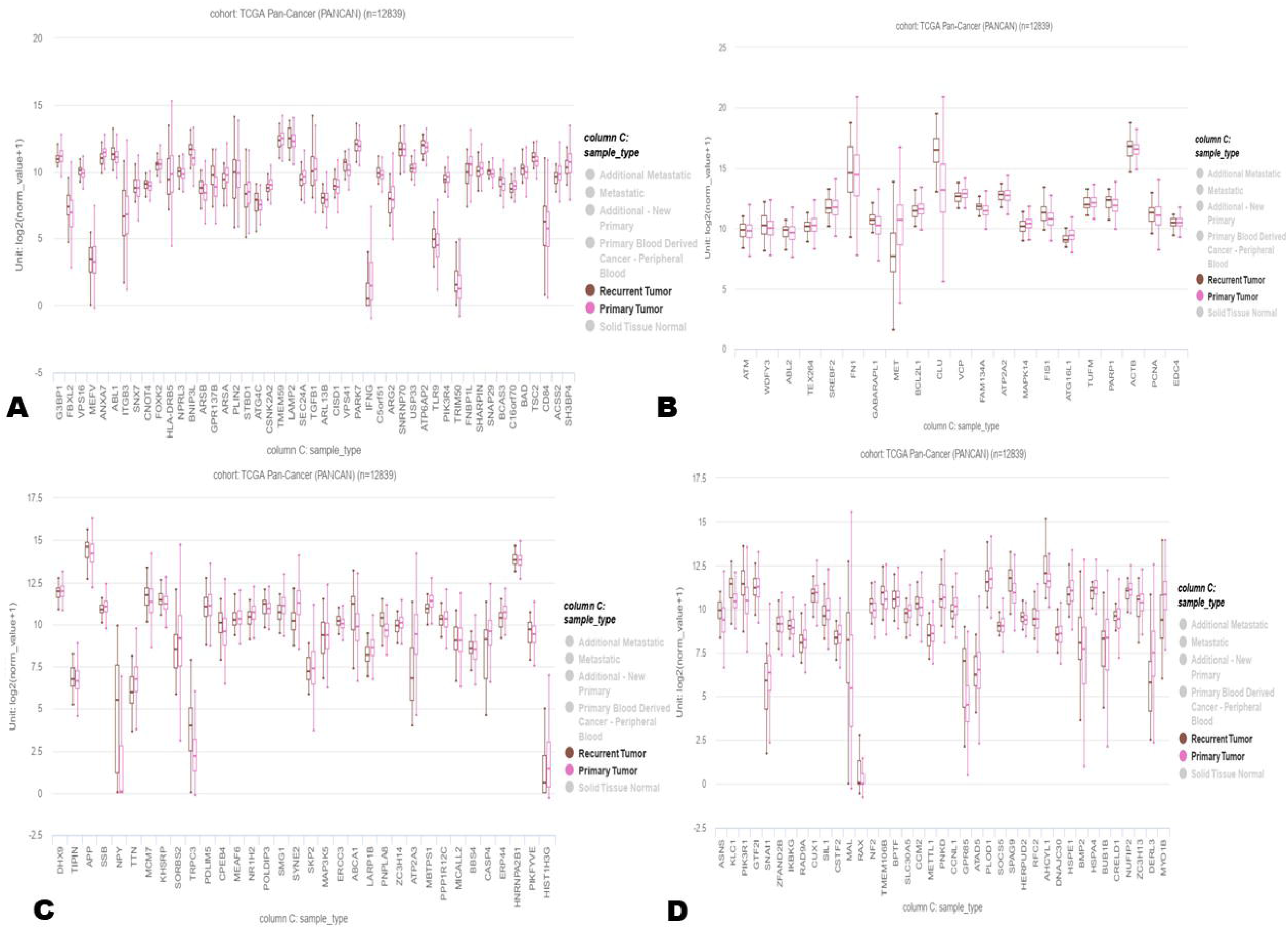
UCSC Xena exploitation for tumour recurrence. Genes related to (A) autophagy (B) mitochondrial elongation & dysregulation (C) & (D) ER stress, short ER fragments & ER stress

The shortlisted genes associated with recurrent tumors were further analyzed for mutation analysis using the GSCA (Gene Set Cancer Analysis) online tool [13]. The results of gene variation were tabulated in Supplementary Information 4. BNIP3L exhibited 12.54% total amplification and 31.53% total deletion, with heterozygous deletion accounting for 29.15%. The pie chart illustrating the mutation distribution for each gene across cervical cancers is presented in Figure 5(A). Figure 5(B) depicts the correlation between genetic variation and gene expression for each gene.

**Figure 5.**
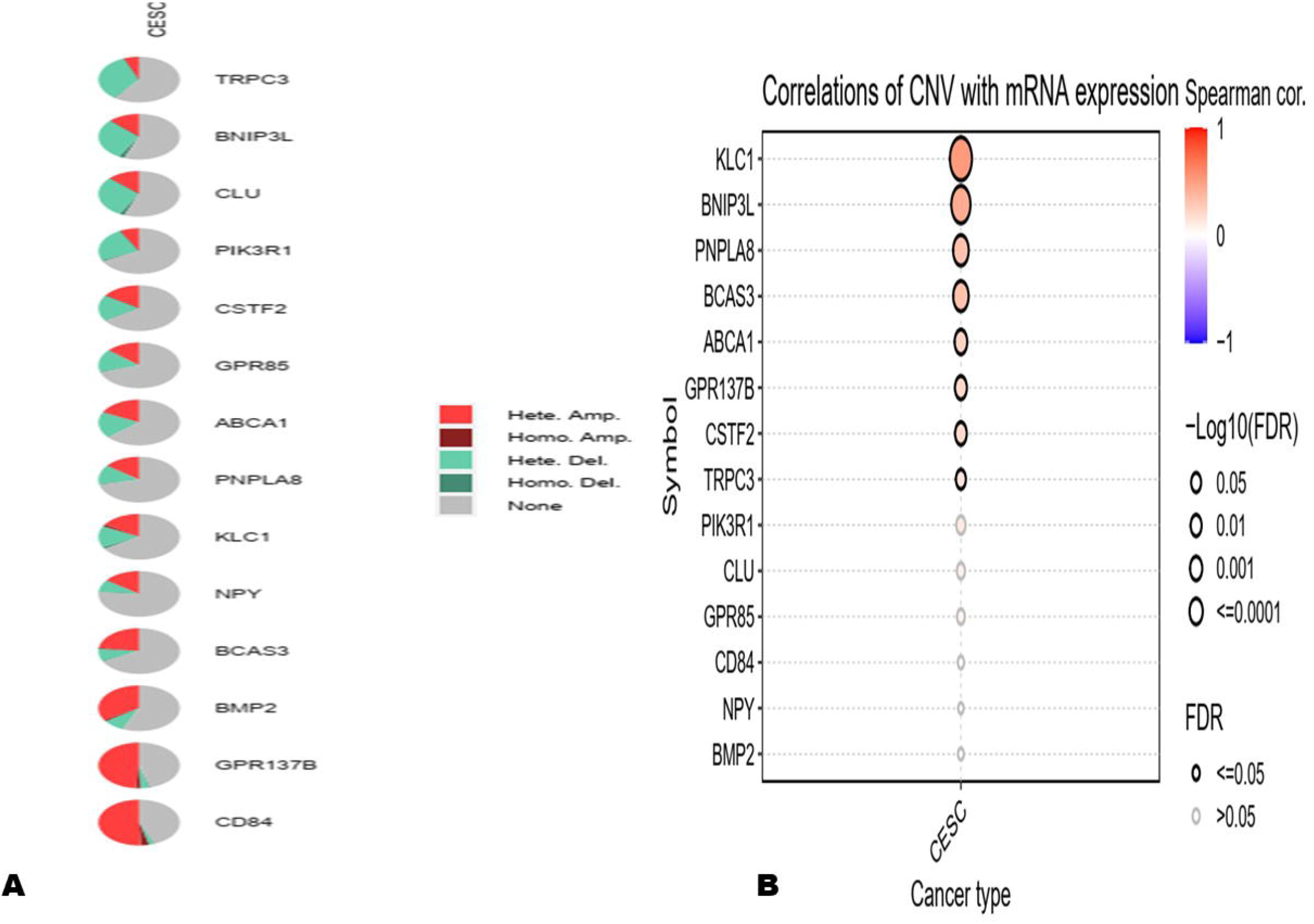
GCSA analysis for genetic variation in tumour recurrence genes (A) amplification percentage in each gene (B) relation of amplification and mRNA expression

### 3.4. BNIP3L is highly expressed in SiHa-Res cells

To confirm that BNIP3L is aberrantly expressed in radiation-resistant cells, SiHaN and SiHa-Res cells were used for expression analysis. Our findings show that BNIP3L is overexpressed in SiHa-Res cells. qPCR, immunofluorescence (IF), and western blot analyses were performed on SiHaN and SiHa-Res cells to investigate BNIP3L expression levels. BNIP3L levels were nearly 2.5-fold higher in SiHa-Res cells compared to SiHaN cells (Figure 6B). Similarly, western blot data revealed that SiHa-Res cells express BNIP3L 1.16-fold more than control SiHaN cells. ROCK2 was used as an indicator of radiation response in both qPCR and western blot analyses [8]. Further to confirm identify the mitochondrial metabolism level and mitochondrial membrane in resistant cells, GAPDH and TOMM20 expression levels were monitores.d metabolism. Our immunofluorescence data on GAPDH and TOMM20 inmonitored with IF. Figure 6(A), indicate GAPDH is downregulated and TOMM20 is upregulated in SiHa Res cells.

**Figure 6.**
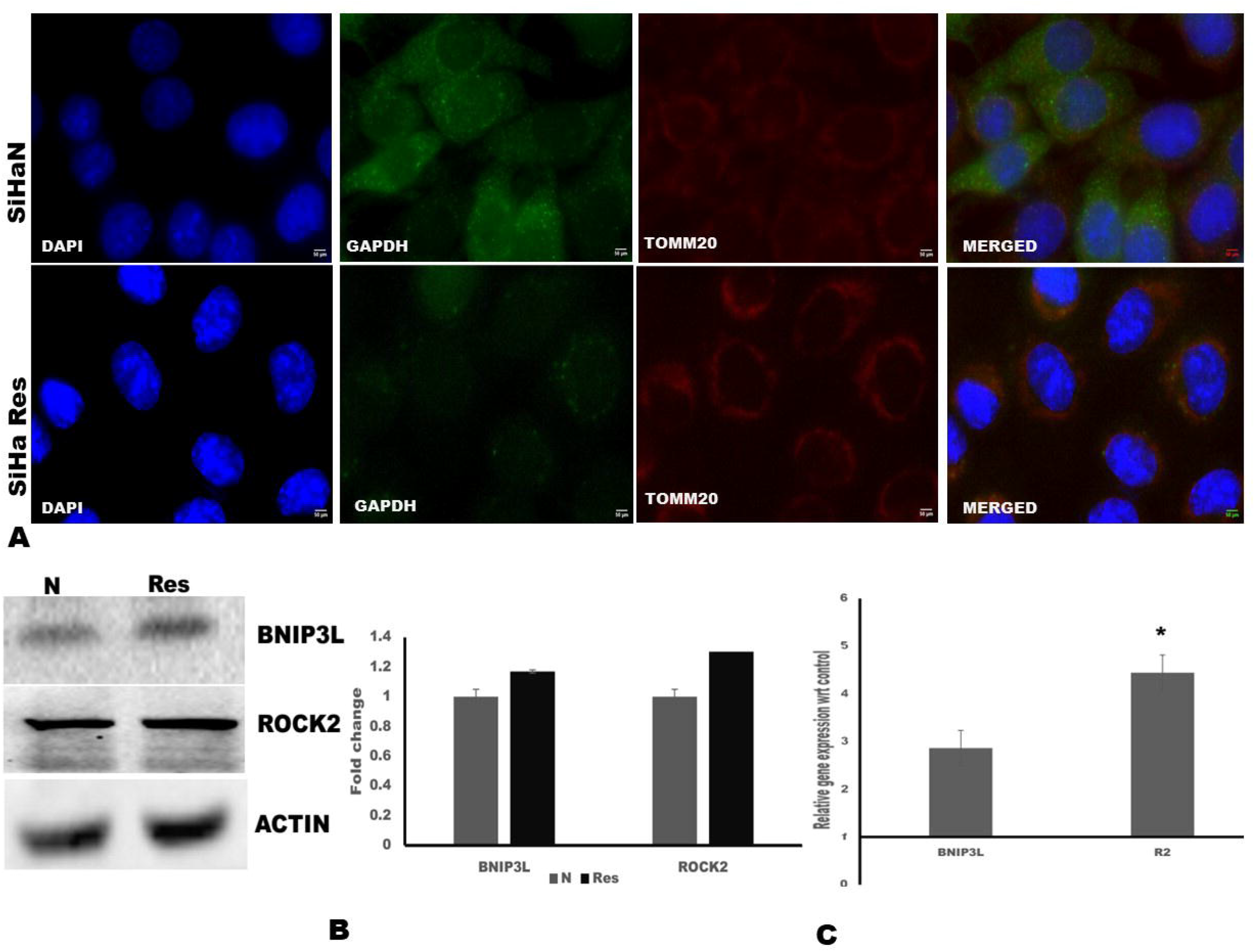
Mitochondrial status in Resistant cells. (A) IF of SiHaN cells and SiHa-Res with GAPDH and TOMM20. SiHa-Res expresses lower GAPDH and higher TOMM20. BNIP3L expression in SiHa-Res cells (B) Western blot and (C) qPCR with BNIP3L and ROCK2 upregulation. * Indicates p< 0.05.

## 4. Discussion

The Mitochondrial Associated Membrane (MAM) serves as a hub for proteins responsible for apoptosis, autophagy, cell proliferation, survival, oncogenes, tumor suppressors, and more. MAM is a dynamic structure that adapts to cellular requirements [7]. Damage induced by radiation therapy on DNA is effectively countered by the DNA repair machinery [14], ultimately leading to radiation resistance. To understand the radiation-induced damage to mitochondria and the endoplasmic reticulum, as well as their mechanisms for survival, we developed radiation-resistant cells. Radiation resistant cells had enhanced survival and clone formation compared to normal SiHa cells, whereas its proliferation is reduced.

The MAM structure of SiHa-Res was studied using TEM. SiHa-Res cells adapt to radiation-induced stress by forming lipid droplets, elongation of mitochondria, induced mitochondria autophagy or mitophagy, and shortened RER, which were absent in SiHaN cells. Yang T. et al. reported a correlation between the number of lipid droplets and radiation dose in radiation-resistant lung cancer cells compared to primary lung cancer cells [15]. Lipid droplets are the main source of energy for the cell during cancer cell invasion. The density and motility of lipid droplets correlate with the invasiveness of cancer and are closely associated with chemoresistance [16], metastasis, and resistance to cell death *in vivo* [17]. The biogenesis of lipid droplets in a cell is a response to nutrient deprivation, which is a stress response. Lipid droplets protect cellular integrity, maintain membrane and organelle homeostasis, and regulate autophagy [18]. The presence of lipid droplets in SiHa-Res cells could be a factor contributing to radiation-induced resistance. The presence of short fragments of RER in SiHa-Res cells, could be an indicator of ER stress [19] or changes in the ER associated with the emergence of lipid droplets [20].

Mitochondrial elongation and mitophagy were observed in SiHa-Res cells. Mitophagy can act as a double-edged sword in radiation sensitivity; it could either induce radiation resistance or sensitize cells to radiation [21]. SiHa-Res cells may have adapted mitophagy as a mechanism of radiation resistance. Mitochondrial elongation represents an adaptation of mitochondria through mitochondrial fusion to evade mitophagy or autophagy [22]. The elongation of mitochondria in SiHa-Res cells suggests the potential for cells to survive in a radiation-induced resistance state.

MFN2, a protein found in the mitochondrial outer membrane, is involved in ER-Mito tethering, mitochondrial fusion and fission, lipid droplet formation, mitochondrial dysfunction, mitophagy, etc. [23, 24]. MFN2 dysfunction or variation leads to neuropathological diseases. The major changes resulting from the MFN2 variation are alterations in the interaction sites between the endoplasmic reticulum (ER) and mitochondria [10]. The downregulation of MFN2 indicates either the disruption of MAM structure or mitochondrial dysregulation. A study on mTORC2 suppression and its impact on MFN2 supports MFN2 downregulation is responsible for cancer progression in breast cancer [25]. Conversely, in ovarian cells, the downregulation of MFN2 sensitizes resistant cells to cisplatin [26]. Further studies are required to understand the exact role of MFN2 in radiation-resistant SiHa cells.

Tumor recurrence is mainly attributed to chemotherapy failure. Therapy or drug resistance is often caused by gene alterations such as mutations or amplifications [27]. Therefore, *the in-silico* analysis was mainly focused on tumour recurrence. BNIP3L is found to be one of the genes expressed in recurrent tumours. BNIP3L, present in the mitochondrial outer membrane, acts as a receptor for mitophagy or mitochondrial autophagy [28] and plays a significant role in cancer recurrence. Gene amplification analysis indicates that BNIP3L has heterozygous amplification, underscoring its importance as a therapeutic target. Hence, the expression of BNIP3L in SiHa-Res cells was validated using qPCR and Western blot. BNIP3L is highly expressed in SiHa-Res cells according to our western blot and qPCR data. A study by Yan C et.al. confirms BNIP3L acts as mitophagy regulator and its inhibition sensitize colorectal cancers to Doxorubicin [29]. Further studies are required to elucidate the signaling cascade of BNIP3L related to radiation resistance in cervical cancers.

## 5. Conclusion

The Mitochondrial Associated Membrane (MAM) is a dynamic structure that varies according to cellular requirements. The interaction between the endoplasmic reticulum (ER) and mitochondria was disrupted in radiation resistance, indicating damage induced in both organelles. However, these stresses or damages were overcome by the adaptability of these organelles, leading to cell survival under stress conditions. This work underscores the importance of MFN2 and BNIP3L in radiation resistance. Future research could focus on identifying the signaling mechanisms of these proteins in radiation resistance.

## ^1^Abbreviation

MAM: Mitochondrial associated membrane
SiHaN: Untreated SiHa cells
SiHa-Res: Radiation resistant SiHa cells
LINAC: Linear particle accelerator

## Acknowledgements

This work was funded under ICMR DHR women scientist scheme. (Sanction no. R.12013/17/2021-HR/E-office 8113319). We acknowledge the help of radiation faculty and staffs of St. Johns medical college hospital for irradiating the samples. We also thank Mr. Sathish, Dept. of neuropathology, NIMHANS for his support in processing TEM samples.

